# RecA balances genomic stability and evolution using many successive mismatch tolerant homology tests

**DOI:** 10.1101/2022.12.09.519633

**Authors:** Mara Prentiss, John Wang, Jonathan Fu, Chantal Prévost, Veronica Godoy-Carter, Nancy Kleckner, Claudia Danilowicz

## Abstract

A double-strand break (DSB) must usually be repaired with as little alteration to the genome as possible, though some rare alterations provide valuable genomic evolution. In *E*.*coli*, a DSB undergoes resection to give 3’ ssDNA tails. These invading strand tails are loaded with RecA protein and then rapidly search the genome for the corresponding (allelic) partner. Thus, a searching ssDNA/RecA filament must almost never make stable non-allelic contact; therefore, it has been puzzling that RecA forms stable products that join partially homologous sequences. Homology testing by RecA family proteins begins with an 8-bp test, followed by successive homology tests of base pair triplets. Here we introduce a highly simplified homology recognition model to highlight how mismatch sensitivity could affect non-allelic pairing in bacterial genomes. The model predicts that even if each triplet test accepts 2 mismatches, RecA can have ∼ 95% probability of establishing allelic pairing after a DSB in *E. coli*; however, that accuracy requires homology testing ⪆50 contiguous base pairs, consistent with the homology lengths probed *in vivo*. In contrast, if no mismatches are accepted testing 14 bp is sufficient, and testing more base pairs does not reduce non-allelic pairing because bacterial genomes contain long repeats.

## Introduction

In this work we consider the known stages of RecA mediated homologous recombination in the context of the sequences of bacterial genomes. To study how accepting mismatches in heteroduplex products might influence DSB repair accuracy *in vivo*, we developed a highly simplified model of homology recognition (see Methods and Materials). The model is informed by experimental results, simulations, modeling, and *in vitro* experiments (Danilowicz et al., 2015, Sagi et al., 2006, Yang et al., 2015, Bazemore et al., 1997, Cox, 2007, Cox and Lehman, 1981, Lin et al., 2019, Qi et al., 2015, Hsieh et al., 1992, Prentiss et al., 2015, Lee et al., 2015). The model is designed to demonstrate how effectively a highly mismatch tolerant RecA could target allelic pairing.

In the highly simplified model, if a homology test is failed, the invading strand completely separates from the dsDNA and homology testing begins again with the invading strand paired to a different position in the dsDNA. In contrast, if a homology test is passed the heteroduplex formed by pairing an invading strand tail to a complementary region in an unbroken chromosome is extended through the bases that passed the homology test. Homology testing begins with an 8 bp test (Qi et al., 2015, Danilowicz et al., 2015) that accepts one mismatch (Danilowicz et al., 2015). The vast majority of incorrect pairings are rapidly rejected by that 8 bp test. In the rare cases that pass the 8 bp test, the heteroduplex extends over those 8 bp, and homology testing progresses through successive base pair triplets (Lee et al., 2015). In the highly simplified model, a triplet passes a homology test if the number of mismatches in the triplet is ≤ N_mismatch_. *In vivo*, ∼50 contiguous homologous bp (Watt et al., 1985, Lovett et al., 2002) are required to give a stable product at a reasonable frequency, suggesting that *in vivo* more than 50 contiguous base pairs are usually homology tested. We define L_test_ as the number of bases that are homology tested before forming an irreversible final heteroduplex product. To quantify the accuracy of final heteroduplex products, we define stringency as the fraction of products that reach L_test_ and join corresponding sequence regions in the invading and complementary strands. We then explore how stringency is influenced by L_test_ and N_mismatch_ and study how long repeats in bacterial genomes affect RecA mediated homologous recombination.

## Results and Discussion

### Highly simplified model results for random genomes

First, we consider the probability that one side of a double strand break in a 5 Mbp sequence consisting of randomly chosen bases (random genome) will lead to an incorrect final heteroduplex product as a function of L_test_. If the homology tests accept no mismatches, no accepted heteroduplex product will contain a mismatch; however, perfectly sequence matched heteroduplex products can still be incorrect if they join different copies of repeats. Figure 1 shows that for a random genome, if no mismatches are allowed in any homology test and L_test_ = 14, then < 2% of final heteroduplex products would be incorrect (equation 5).

**Figure 1:**
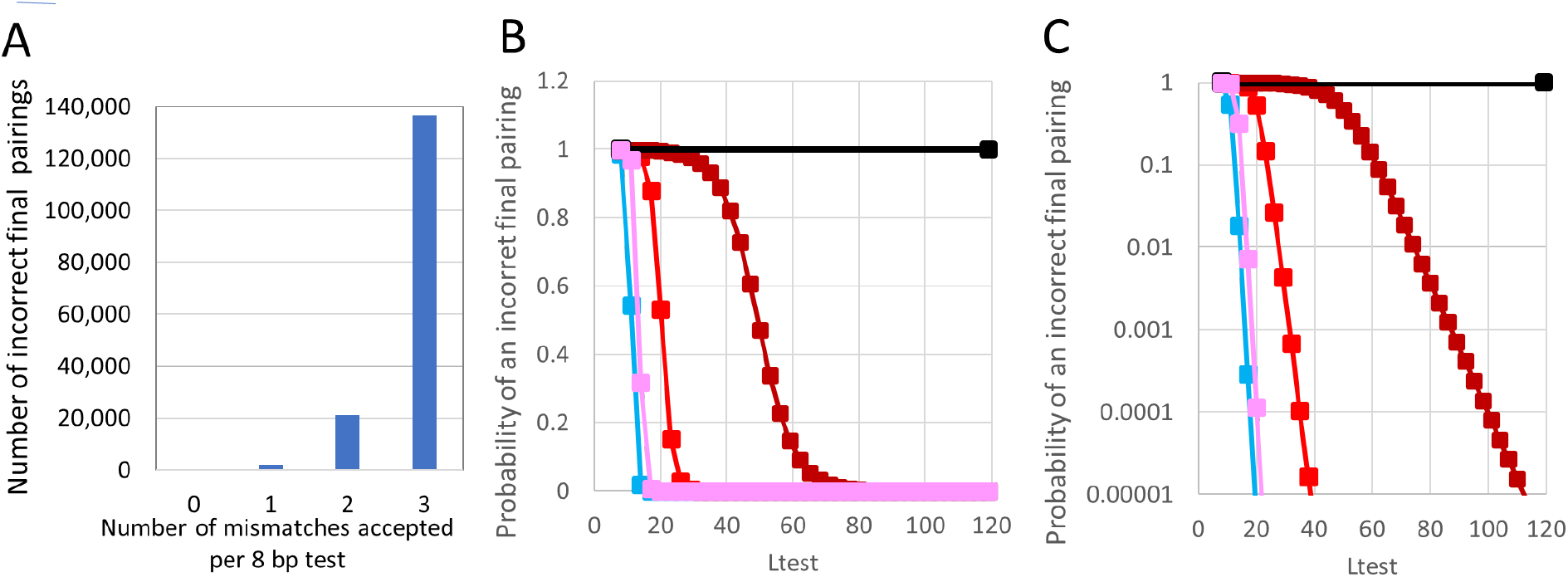
Probability that after a DSB a mistaken pairing in a random 5 Mbp genome will pass various homology tests. A. Number of possible incorrect pairings that would pass the initial 8 bp test vs. the number of mismatches accepted by the 8 bp test. B. Probability that when the heteroduplex length first reaches L_test_ the heteroduplex is incorrect vs. L_test_ when no mismatches are accepted in any homology test (blue). The results for homology tests that accept 1 mismatch in the 8 bp test and 0 (pink), 1 (red), 2 (dark red), or 3 (black) mismatches in each triplet test (N_mismatch_ = 0, 1, 2, or 3) are also shown. C. Same as B but with a logarithmic scale on the y-axis.

In contrast, when homology tests accept mismatches some final heteroduplex products can contain mismatches. Figure 1 illustrates results of the highly simplified homology testing model when N_mismatch_ = 0 bp (pink), N_mismatch_ = 1 bp (red), or N_mismatch_ = 2 bp (dark red). Unsurprisingly, for a given L_test_, increasing the number of accepted mismatches decreases the stringency; however, when N_mismatch_ < 3,stringency always increases with L_test_, and stringency > 10^−7^ can be obtained by choosing L_test_ > 150 bp. Importantly, the stringency for RecA’s iterative testing is much better than the stringency that would result if all L_test_ base pairs were tested simultaneously and the total number of accepted mismatches is the same as the total number that could be accepted by RecA when 0 < N_mismatch_ < 3 (Figure 1 — figure supplement 1).

**Table 1:** Probability that after a DSB a mistaken pairing will pass various homology tests in a random 5 Mbp genome.

| Test size | Number of accepted mismatches per 8 bp test | % Probability of Passing an 8 bp test | Number of incorrect pairings after 8 bp test | % Probability passed pairing is wrong | Test size | Nmismatch = Number of accepted mismatches per triplet | % Probability of passing 1 triplet test | % Probability of passing 3 triplet tests | % Probability of passing 20 triplet tests | % Probability passed pairing is wrong after 8 bp + 3 triplets | % Probability passed pairing is wrong after 8 bp + 20 triplets |
| --- | --- | --- | --- | --- | --- | --- | --- | --- | --- | --- | --- |
| 8 bp | 0 | 0.002 | 76 | 98.706 | 3 bp | 0 | 1.6 | 0.000 | 7.5E-35 | 0.04 | 0.0000000000 |
| 8 bp | 1 | 0.038 | 1907 | 99.948 | 3 bp | 1 | 15.6 | 0.380 | 7.3E-15 | 87.87 | 0.0000000000 |
| 8 bp | 2 | 0.423 | 21133 | 99.995 | 3 bp | 2 | 57.8 | 19.310 | 2.3E-3 | 99.73 | 3.1979503379 |
| 8 bp | 3 | 2.730 | 136490 | 99.999 | 3 bp | 3 | 100.0 | 100.000 | 100 | 99.95 | 99.9475987109 |

### Highly simplified model results for bacterial genomes

Previous work that has considered the mismatch tolerance of RecA (Hsieh et al., 1992, Lee et al., 2006, Xiao et al., 2006, Bazemore et al., 1997, Sagi et al., 2006, Danilowicz et al., 2015, Rosselli and Stasiak, 1991, Qi et al., 2015, Savir and Tlusty, 2010) has not considered how that mismatch tolerance could affect allelic pairing in bacterial genomes. In Figure 2, the blue, pink, bright red, dark red, and black curves with the hollow square symbols and dotted lines are the same as those shown by the solid squares and lines in Figure 1 C. Those results for homology testing a random genome can be compared to the result for homology testing an *E*.*coli* MG1655 genome, which are indicated in Figure 2 by the solid lines with the triangular symbols. At small L_test_ values the results for the random genome and the bacterial genome are indistinguishable, but at larger L_test_ values the results radically depart.

**Figure 2:**
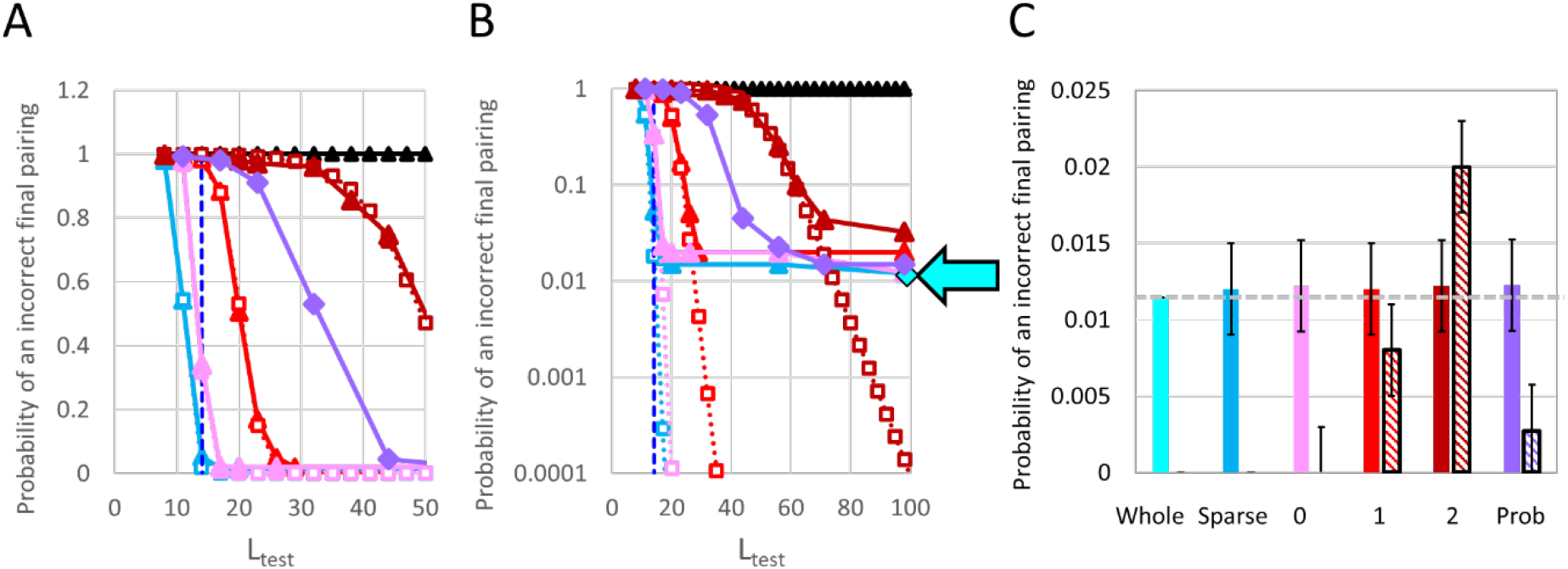
The probability that after a DSB RecA will create a final incorrect pairing as a function of L_test_ for the random genome (square symbols) and the *E.coli* MG1655 genome (triangular symbols). A. The light blue lines show results for sparse homology testing that requires complete sequence matching. The pink, bright red, dark red, and black lines indicate results for sparse homology testing when the 8 bp test that accepts one mismatch is followed by triplet tests that accept N_mismatch_ = 0, 1, 2, and 3 mismatches, respectively. The purple curve shows the results for a probabilistic model in which the probability of passing a triplet homology test is 100%, 75%, 50%, and 25% for 0, 1, 2, and 3 mismatches, respectively, and the 8 bp test has 100% and 25% chance of accepting 0 or 1 mismatch, respectively. The dark blue vertical dashed line indicates L_test_ = 14. B. Same as A with a logarithmic y-axis, but without the line indicating L_test_ = 14 and with the cyan diamond highlighted by the cyan arrow which is the exact result for the complete genome. C. The solid and hashed bars show the fraction of final pairings that join different copies of exact repeats and the fraction that contain mismatches, respectively. The color of the bar corresponds to the homology test represented. The color code is the same as in the rest of the figure. The fraction of DSB that leads to final pairing of exact repeats is very similar for all homology testing systems, but the number of final pairings that contain mismatches depends on the details of the homology testing system. The error bars indicate the rms variation between different trials.

The vertical dashed line at L_test_ = 14 indicates the approximate position at which stringency vs. L_test_ saturates for the *E. coli* MG1655 genome when homology testing rejects all mismatches. The saturation value of ∼ 1% is obtained for both sparse sampling and sampling of the entire genome (Figure 2 —figure supplements 1-4). The saturation results from non-allelic sequence matched pairings between different copies of long repeated sequences in *E. coli* MG1655. Importantly, other bacteria also contain long repeats (Figure 2 — figure supplements 2-4) that would make stringency saturate with increasing L_test_. In sum, for bacterial genomes stringency as a function of L_test_ saturates at ∼1% because incorrect final pairings join different copies of long repeated sequences, and the L_test_ value required to reach saturation increases with N_mismatch_.

### *In vitro* experiments

*In vivo*, the progress of strand exchange is almost certainly probabilistic. Thus, to gain insight into the probability that strand exchange progresses through mismatched regions into an adjacent sequence matched region we performed *in vitro* experiments in which mismatched regions were surrounded by homologous regions (Figure 3 —figure supplement 1). We first considered strand exchange interactions with evenly spaced mismatches, as shown in Figure 3. The results indicate that strand exchange interactions with 1 mismatch per 3 bp do not form significant strand exchange products even if the evenly spaced mismatches are flanked on the 5′ side by > 8 contiguous homologous base pairs that could pass the initial 8 bp homology test. In contrast, when the invading strand has 1 mismatch per 6 bp it can pass the initial 8 bp test and form readily observable strand exchange products, consistent with *in vivo* results in yeast (Anand et al., 2017). Experiments also indicate that strand exchange can progress through 6 contiguous mismatched base pairs (Figure 3 — figure supplement 1), which is also consistent with results for the eukaryotic analog Rad51 (Ristic et al., 2010, Holmes et al., 2001). In sum, triplet homology testing by RecA family proteins is both more and less stringent than predictions of the highly simplified model for N_mismatch_ = 1 since interactions with 1 mismatch per triplet are often rejected and RecA mediated strand exchange can progress through 6 contiguous mismatched bases.

**Figure 3.**
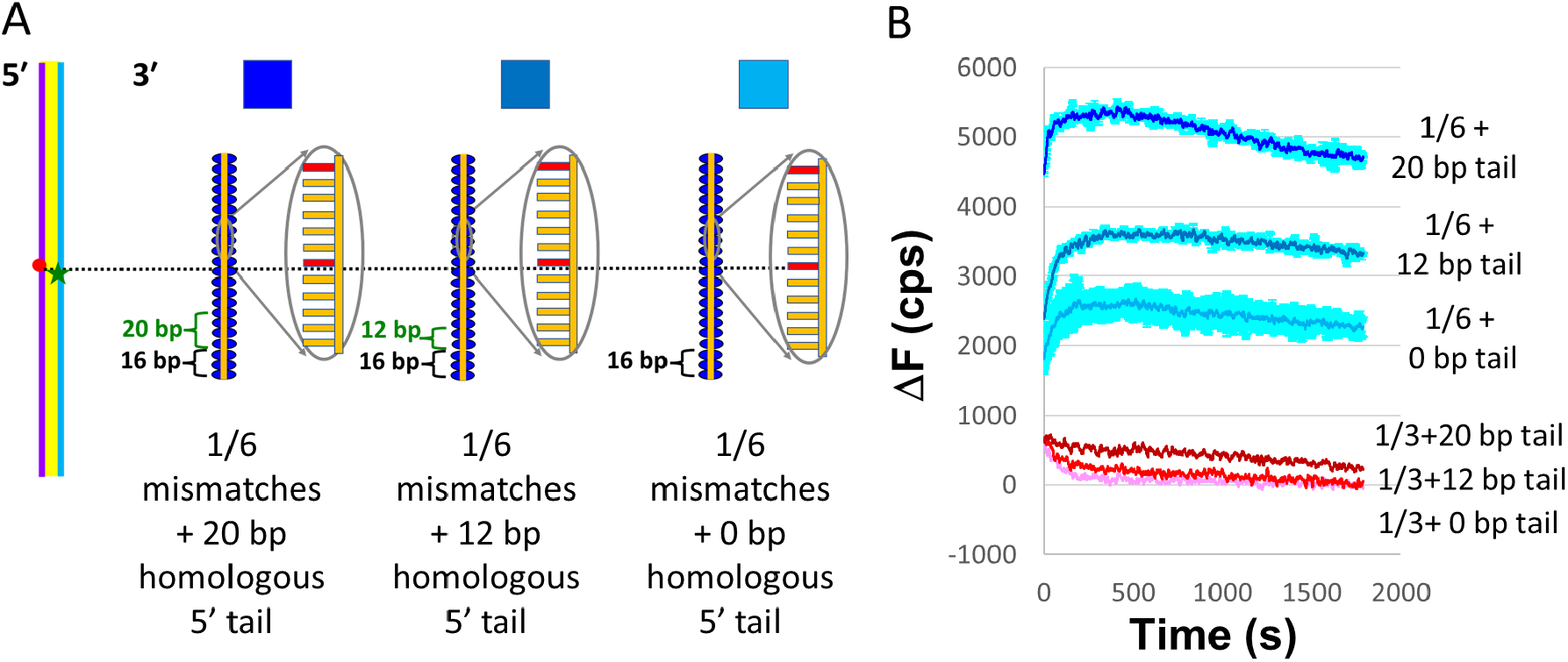
Strand exchange monitored using FRET. A. Schematics of interactions of 180 bp dsDNA with 98 nt filaments containing different degrees of homology. The purple, light blue, and orange lines represent the complementary, outgoing, and invading strands. The dark blue ovals represent RecA. The red circle (rhodamine)and green star (fluorescein) show the locations of the fluorophores along the 180 bp dsDNA. All of the invading strands contain 98 nt. The 16 nt nearest the 5′ end of the invading strand are indicated by the black brackets and are always heterologous to the dsDNA. The green brackets indicate regions of the invading strand that are completely homologous to the dsDNA. The orange regions of the invading strand are homologous except for periodic mismatches that are indicated by the red bars. 1/6 mismatches are shown. B. Change in fluorescein emission (ΔF) vs. time curves. The dark blue, medium blue, and light blue curves indicate results for 1/6 periodically spaced mismatches with a 20 bp, 12 bp, or 0 bp homologous tail on the 5′ side. The dark red, red, and pink curves indicate results for 1/3 periodically spaced mismatches with a 20 bp, 12 bp, or 0 bp homologous tail on the 5′ side.

### More realistic modeling

The experimental results on the mismatch sensitivity of strand exchange led us to consider some simple probabilistic models of how strand exchange progresses through triplets (purple lines in Figure 2, Figure 2 — figure supplements 5 and 6). Importantly for all of the probabilistic models we considered, stringency saturates at L_test_ ∼ 50-75 bp. Furthermore, even though the probabilistic models sometimes accept triplets with 3 mismatches, if the probabilistic model also sometimes rejects triplets with 1 mismatch, then when L_test_ = 98 the fraction of final products that contains mismatches can be smaller than the fraction predicted by the highly simplified model with N_mismatch_ = 1 (Figure 2C).

### Ideas and Speculation

Importantly, if N_mismatch_ > 0 and L_test_ is small, almost all final pairings include mismatches (Figures 1 and 2, Figure 2 — figure supplements 5 and 6). Small L_test_ values can arise if strand exchange starts close to the 3′ end of the invading strand. Thus, *in vivo* if small L_test_ values often create irreversible products, then genomic rearrangement would also be quite frequent; however, irreversible pairing may depend on DNA polymerization by Pol IV (Henrikus et al., 2018b, Henrikus et al., 2018a). Importantly, *in vitro* results suggest that extended DNA polymerization by Pol IV very rarely occurs unless the heteroduplex length, L, exceeds ∼ 50 bp (Lu et al., 2019).Thus, Pol IV polymerization may play an important role in rejecting short incorrect heteroduplex products formed when L_test_ is small.

Even if L = L_test_ = 98 bp, approximately 1% of final pairings join different copies of exact repeats. Those non-allelic pairings would allow extensive Pol IV polymerization and would not be reversed by MutS; therefore, those pairings between long repeats could lead to genomic rearrangement if they are resolved by a crossover. In addition, when L = L_test_ = 98 bp another ∼ 1% of the final pairings are non-allelic but could be reversed by MutS because they do contain mismatches (Worth et al., 1994). Furthermore, we note that any sequence testing system can trade-off mismatch tolerance against the number of base pairs that are homology tested.

Searching speed is an important requirement for many sequence testing systems. Previous work suggested that features that make strand exchange mismatch tolerant also make strand exchange more free-energetically favorable (Danilowicz et al., 2015, Yang et al., 2015). That favorable free energy may allow strand exchange to progress more quickly. Additionally, we speculate that highly accurate sequence recognition requires rigidly held Watson-Crick partners that would slow homology testing. Thus, we propose that RecA homology testing is mismatch tolerant for the following reasons: 1. mismatch tolerance is a consequence of structural features that speed DSB repair; 2. mismatch tolerance allows strand exchange between allelic partners that include mismatches; 3. mismatch tolerance permits some desirable non-allelic pairings.

In sum, the level of genomic alterations produced during recombinational repair reflects a critical balance between the rate of highly mismatch tolerant RecA-mediated strand exchange and the rate of intervention by other cellular components. The balance is presumably tuned by evolutionary forces to meet the requirements of rapid repair, genetic stability, and genetic variation, which may vary according to the cellular environment.

## Acknowledgements

We acknowledge useful interactions with Benjamin Tang and funding from the Chu Family Foundation and Harvard University.

## Materials and Methods

### Simplified model of double strand break repair that follows the RecBCD pathway

Having strand exchange depend dominantly on the kinetics of heteroduplex extension rather than the mismatch dependence of heteroduplex lifetimes was previously suggested (Sagi et al., 2006) and is supported by modeling and experiments indicating that the stability of heteroduplex products is insensitive to mismatches (Bazemore et al., 1997, Danilowicz et al., 2015)and by experiments showing that mismatches have a stronger effect on strand exchange than on the stability of heteroduplex product (Danilowicz et al., 2015).

Thus, consistent with experimental results, modeling, and simulations we developed a simplified model of DSB repair that follows the RecBCD pathway. The model has the following features: 1. Homology testing and strand exchange progress in the 5′ to 3′ direction with respect to the invading strand (Cox, 2007, Lin et al., 2019) (Figure 1). 2. The initial homology test considers 8 RecA bound base pairs (Qi et al., 2015, Danilowicz et al., 2015, Hsieh et al., 1992, Yang et al., 2015, Prentiss et al., 2015) and accepts one mismatch(Danilowicz et al., 2015). 3. All subsequent tests consider successive RecA bound base pair triplets (Lee et al., 2015). 4. A triplet can only attempt strand exchange if it is bound to RecA and the RecA bound triplet on its 5′ side has already undergone strand exchange (Yang et al., 2015). 5. Reversal of the heteroduplex product does not proceed backwards in the 3′ to 5′ direction with respect to the invading strand (Danilowicz et al., 2021). 6. The reversal rate of the heteroduplex product is independent of the length of the heteroduplex and the number of mismatches already incorporated in the heteroduplex. 7. After a heteroduplex completely reverses, a new homology test begins with a different registration between the ssDNA-RecA filament and the dsDNA. 8. The registration between the invading and complementary strands becomes irreversible when the heteroduplex length is equal to L_test_, the separation between the 5′ side of the heteroduplex and the 3′ end of the invading strand since reaching the 3′ end allows DNA polymerization that makes the registration between the complementary and invading strands irreversible (Lu et al., 2019). In the simplified model the pairing between the invading and complementary strands becomes final the first time that the length of the heteroduplex is equal to L_test._

In reality, strand exchange through a mismatched triplet may be attempted many times before the complementary and invading strands completely separate. Complete separation only occurs when the heteroduplex on the 5′ side of the triplet entirely reverses. Thus, each step in the simplified model does not represent a single attempt to extend the heteroduplex into the mismatched triplet, each step in the simplified model considers whether one of those attempts succeeds before the heteroduplex on the 5′ side of the triplet completely reverses.

We note that in our probabilistic models the probabilities chosen for the incorporation of mismatched triplets were somewhat arbitrary. They were merely designed to illustrate general effects of extending the highly simplified model. One could try to assign a binding energy to a perfectly matched triplet, and a binding energy penalty for each mismatch within a triplet; however, previous work has shown that a single mismatch in 6 base pair B-form DNA can result in a collective effect that destabilizes all size bases (Cisse et al., 2012). Since heteroduplex products consist of nearly B-form triplets (Chen et al., 2008), a single mismatch is likely to have a collective effect that destabilizes the triplet. The destabilizing effect may also depend on the position of the mismatch in the triplet and the exact sequence of the triplet. Given all of these points, we did not feel justified in assuming that the binding energy for a mismatch triplet depends linearly on the number of sequence-matched base pairs in the triplet. Furthermore, the positive charges near the primary binding site in RecA (Yang et al., 2015) and the RecA residues intercalated between the rises in the base pair triplets (Chen et al., 2008, Yang et al., 2015) imply that previous measurements of the free energy penalty for mismatches in B-form dsDNA are unlikely to apply to heteroduplex dsDNA bound to the primary binding site in RecA. Finally, both theory and experiment suggest that dsDNA bound to site I is insensitive to mismatches. (Danilowicz et al., 2015). In sum, given all of the uncertainties regarding the free energy penalty due to mismatches in heteroduplex triplets bound to the primary binding site in RecA, we explored a range of probabilities of progressing through mismatched triplets to determine the sensitivity of the results to the probabilities.

### Probabilities of incorrect pairings for a homology search following a DSB in a random genome

For a random genome, the probability that one randomly chosen base will match another randomly chosen base is p = ¼. The probability of finding m or more correctly paired bases in a randomly chosen sample of n base pairs is then

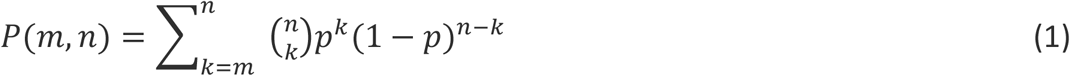

For the 8 bp test, n = 8 and for the triplet tests n = 3. In either case m=n-N_mismatch_.

Thus, for the 8 bp test with one mismatch n = 8 and m = 7, and for a triplet test that accepts two mismatches n = 3 and m = 1.

The probability of passing a series of tests is just the product of the probabilities for all of the tests. Thus, for a given L_test_, the probability of passing an 8 bp test followed by a series of triplet tests is the product of the probability of passing the first 8 bp test times the probability of passing (L_test_-8)/3 triplet tests.

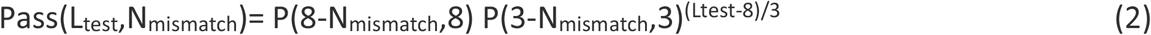

The formula implies that the results for a series of triplet tests that accept only one mismatch per triplet are much more stringent than the results of a test that accepts 1/3 mismatches in L_test_-8 bases.

For a genome that includes L_genome_ randomly chosen bases, and a randomly chosen sequence of length L_test_, in the genome there are on average Pass(L_test_,N_mismatch_) L_genome_ matching sequences of length L_test_.

In DSB repair for a random genome, we assume that the sequence of the searcher is always included in the target; therefore, there is also always one correct pairing between the searcher and the target, regardless of the probability that a randomly chosen searching sequence would find a match. Thus, for a random genome during the homology search that follows a DSB, the average number of pairings that would pass the RecA homology test is:

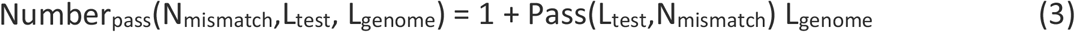

Of those that pass, probability of being correct is then 1 out of the total number or

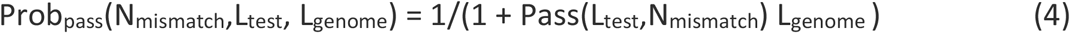

The probability that a pairing passes and is incorrect isthen:

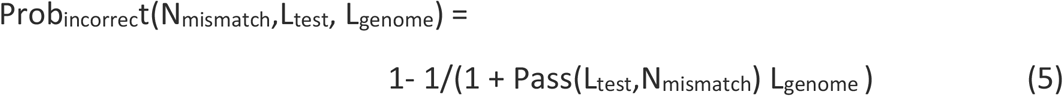

### Probabilities of incorrect pairings for a bacterial genome using the simplified model that is applied sparsely to the sequences of bacterial genomes

A position in the given strand of genome was randomly chosen as the position of the 5′ end of the invading strand. A second position in the given strand of the genome was randomly chosen to represent the testing position in an unbroken chromosome that pairs with the 5′ end of the invading strand. The sequences of the 8 bp starting at the 5′ end and extending in the 3′ direction were then compared. If the number of mismatches was > N_mismatch_, homology testing at that position terminated and a new testing position was chosen. If the number of mismatches was ≤ N_mismatch_, then the next 3 bases on the 3′ side were homology tested. Homology testing terminates and a new testing position is chosen whenever the number of mismatches in a triplet > N_mismatch_. If the length that has passed the homology tests reaches L_test_, then the pairing is considered irreversible. If the test position and the search position are the same, the irreversible pairing is correct. If the test position and search position are different but there are no mismatches, then the irreversible pairing joins two copies of a repeat with length ≥ L_test_. If the irreversible pairing contains mismatches, the pairing is an error. Typically, 100 -10000 different invading strand sequences were chosen, and the results represent the averages of the results for those 100-10000 invading strand sequences out of ∼ 5 Mbp possible invading strand sequences.

### Probabilities of incorrect pairings when N_mismatch_ = 0 for entire bacterial genomes

Each possible 8 bp sequence was assigned a unique mapping number. The bacterial genome was divided into 8 bp sequences, each of which was assigned the corresponding mapping number. The 8 bp sequences were then sorted according to their mapping number, which grouped the entire genome into distinct 8 bp repeats. The total number of incorrect pairing locations for each member of the group is equal to the number of locations in the group -1. The probability that a DSB will lead to an incorrect pairing is then the sum over all groups of repeats divided by the total number of bases in the genome. We repeated the same procedure for 11 bp sequences. To calculate exact repeats for 14, 17, 22, 33, 44, 55, and 99 we created a list of maps encoding a series of sequences with lengths ≤ 11 bp. For example, for L_test_ = 14 we used an 11 bp map and a 3 bp map, and then grouped starting locations with the same values for both maps. That allowed us to calculate the probability that a DSB in a bacterial genome would lead to an incorrect final pairing if it was subject to a homology test that accepts no mismatches over a length L_test_.

### Calculation of the number of distinct long repeat pairs in bacterial genome

To determine the prevalence of longer repeats, each possible 12 bp sequence was assigned a unique mapping number. The bacterial genome was divided into 12 bp sequences, each of which was assigned the corresponding mapping number. The 12 bp sequences were then sorted according to their mapping number, which grouped the entire genome into distinct 12 bp repeats. Longer repeats were probed by extending the 12 bp sequences within each mapping group and counting only those pairs that had no mismatches over the length L_test_. If the number is non-zero, that implies that there must be some repeats that have lengths ≥ L_test_. If most repeats only occur twice, the ratio of the number of distinct pairs to the length of the genome gives the probability that a DSB will lead to an incorrect pairing of that length.

### Probabilities of incorrect pairings for a bacterial genome using the simplified model with Chi sites that sparsely samples bacterial genomes

A Chi site in the given strand of genome was randomly chosen as the position 8 bp from the 3′ end of the invading strand. The invading strand sequence was then the L_test_ bp on the 5′ side of the 3′ end of the invading strand. The remainder of the homology testing was exactly the same as the testing without Chi sites.

### Simplified Probabilistic model of homology testing in triplets

In the first model, a random number generator creates values 0, 1, 2, and 3 with equal probability. A homology test of a triplet then passes the triplet if the number of mismatches is less than or equal to the value determined by the random number generator. Since 0 mismatches is less than or equal to all of those values, a triplet with 0 mismatches has a 100% chance of passing the homology test. Similarly, 1 mismatch, 2 mismatches, or 3 mismatches have a 75%, 50%, or 25% chance of passing the homology test. In the second model, a random number is chosen. If the random number is less than 0.5, the triplet homology test is passed regardless of the number of mismatches; however, if the random number is less than 0.5, then a triplet without a mismatch will have a 100% probability of passing the homology test, whereas a triplet with 1 mismatch has a 50% chance of passing. Thus, for a triplet the probability that 0, 1, 2, or 3 mismatches will pass the test is 100%, 75%, 50%, and 50%, respectively. Thus, for 0, 1, or 2 mismatches the probability of passing either of the probabilistic models is the same. In contrast, for 3 mismatches the first probabilistic test has a 25% of passing the completely mismatched triplet, whereas the second probabilistic test has a 50 % change of passing the completely mismatched triplet.

