## Supplemental Data 1 for "RecA balances genomic stability and evolution using many successive mismatch tolerant homology tests"

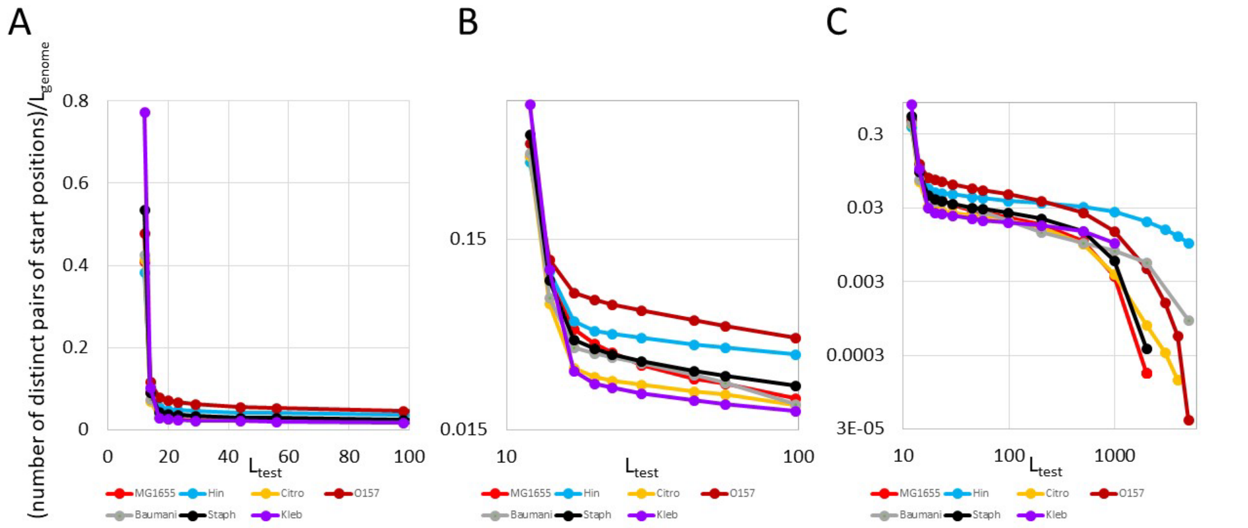


Figure 2 — figure supplement 4 **Ratio of the number of distinct pairs of starting locations in the given strands of bacterial genomes that share a repeat of length L to the genome length as a function of L.** A. The color of the curve corresponds to the genome considered. *E. coli* MG1655, *Haemophilus influenzae* strain NML-Hia-1, *Citrobacter freundii* strain 705SK3, *E. coli* O157:H7 strain JEONG-1266, *Acinetobacter baumannii* strain K09-14, *Staphylococcus aureus* strain Bmb9393, *Klebsiella pneumoniae* strain ATCC BAA-2146 are represented by the red, blue, orange, dark red, gray, black, and purple curves, respectively. Both the x and y axis are linear. B. Same but the x and y-axes are logarithmic. C. Same as B but the x-axis is extended to 6000 bp
