## Supplemental Data 2 for "RecA balances genomic stability and evolution using many successive mismatch tolerant homology tests"

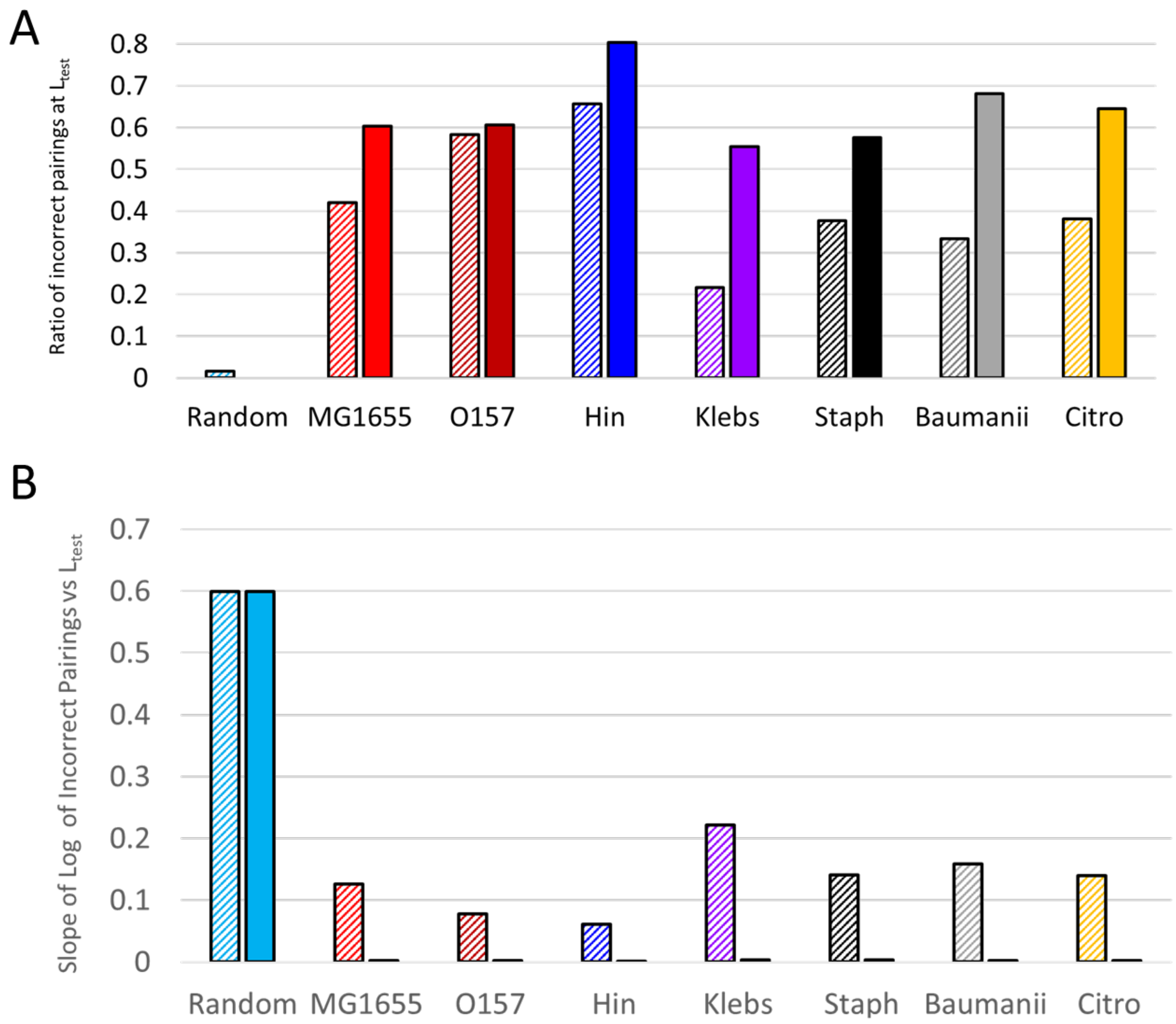


Figure 2 — figure supplement 3 **Saturation in the decrease in incorrect pairings as $L_{test}$ increases. A.** The bars with the striped fills indicate the ratio of the stringency for $L_{test}$ = 17 to the stringency for $L_{test}$ = 14. Smaller bars indicate that stringency increases more quickly with $L_{test}$. The bars with solid fills indicate the ratio of the stringency for $L_{test}$ = 99 to the stringency for $L_{test}$ = 17. For all bacterial genomes, the solid bar is higher than the striped bar. B. Same as A, but the bars represent the absolute value of the slope of the Log of the probability of an incorrect pairing as a function of $L_{test}$. The slopes are always negative. For the random genome, the slopes are always the same for $L_{test} \gtrapprox 14$. In contrast, for the bacterial genomes the slopes from $L_{test}$ = 17 to $L_{test}$ = 99 are barely visible on the graph, indicating that the increase in accuracy with $L_{test}$ strongly saturates when $L_{test} \gtrapprox 17$.
