## Supplemental Data 3 for "RecA balances genomic stability and evolution using many successive mismatch tolerant homology tests"

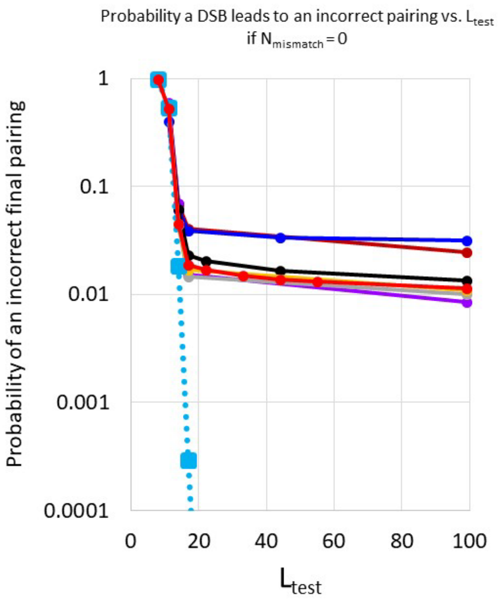


Figure 2 — figure supplement 2 **Probability that a DSB will result in an incorrect final pairing vs. $L_{test}$ if all mismatches are rejected.** A. The dotted blue line represents results for a random genome and are the same as those shown in Figure 1. The colors of the other curves correspond to the genomes considered: *E. coli* MG1655, *Haemophilus influenzae* strain NML-Hia-1, *Citrobacter freundii* strain 705SK3, *Escherichia coli* O157:H7 strain JEONG-1266, *Acinetobacter baumannii* strain K09-14, *Staphylococcus aureus* strain Bmb9393, *Klebsiella pneumoniae* strain ATCC BAA-2146, which are represented by the red, dark blue, orange, dark red, gray, black, and purple curves, respectively.
