## Supplemental Data 4 for "RecA balances genomic stability and evolution using many successive mismatch tolerant homology tests"

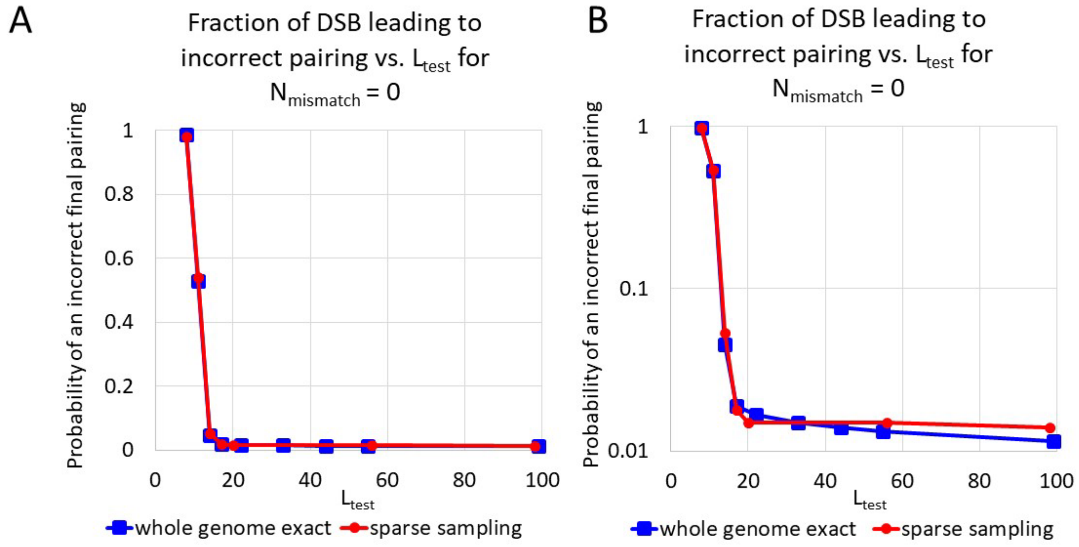


Figure 2 — figure supplement 1 **Predicted incorrect DSB repairs as a function of $L_{test}$ for $N_{mismatch}$ = 0.** A. The red line with circular markers is the same as the red line in Figure 1, which is the result for sparse sampling. The blue line with square markers is the result for complete sampling of the *E.coli* MG1655 genome. Results are very similar. The error in the sparse sampling masks the small increase in stringency that occurs as $L_{test}$ is increased from 17 to 99. B. Same as A but with a logarithmic y axis.
