## Supplemental Data 5 for "RecA balances genomic stability and evolution using many successive mismatch tolerant homology tests"

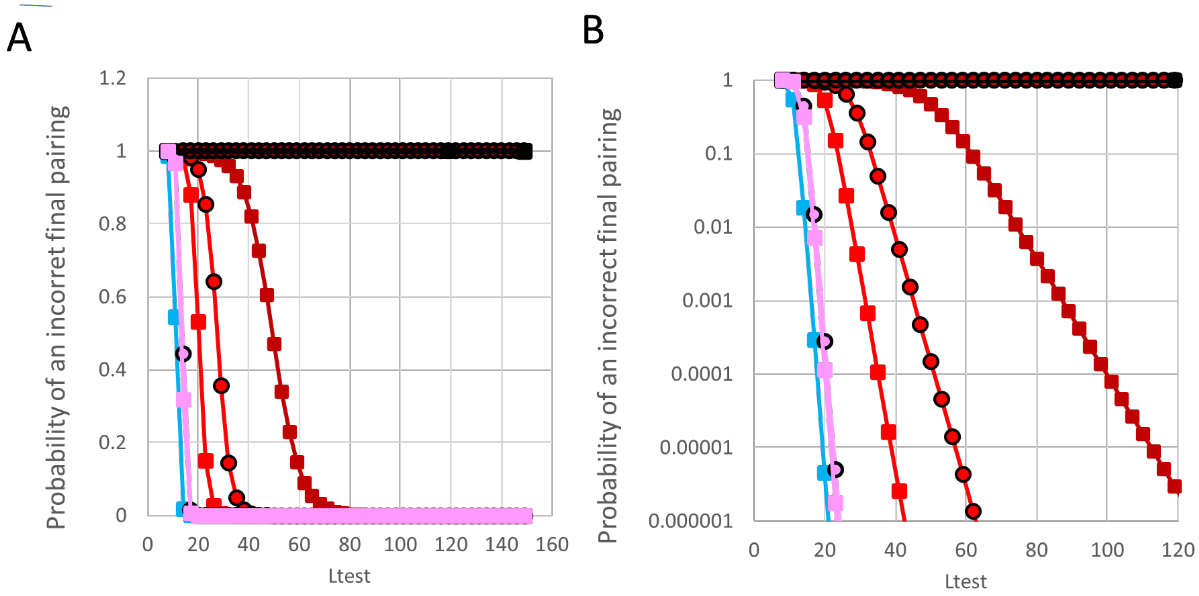


Figure 1 — figure supplement 1. **Probability that when the heteroduplex length first reaches $L_{test}$ the heteroduplex is incorrect vs. $L_{test}$.** A. Results when no mismatches are accepted in any homology test (blue squares) can be compared with results when 1 mismatch is accepted in the 8 bp test and 0 (pink squares), 1 (red squares), 2 (dark red squares), or 3 (black squares) mismatches are accepted in each triplet test ($N_{mismatch}$ = 0, 1, 2, or 3). The circles indicate homology tests that consider all $L_{test}$ bp at the same time and accept as many mismatches as the RecA test that accepts one bp in the initial test and the triplet tests accept $N_{mismatch}$ = 0 (pink circles with black outlines) $N_{mismatch}$ = 1 (red circles with black outlines) or $N_{mismatch}$ = 2 (dark red circles with black outlines). B. Same as A but with a logarithmic scale on the y-axis.
