## Supplemental Data 6 for "RecA balances genomic stability and evolution using many successive mismatch tolerant homology tests"

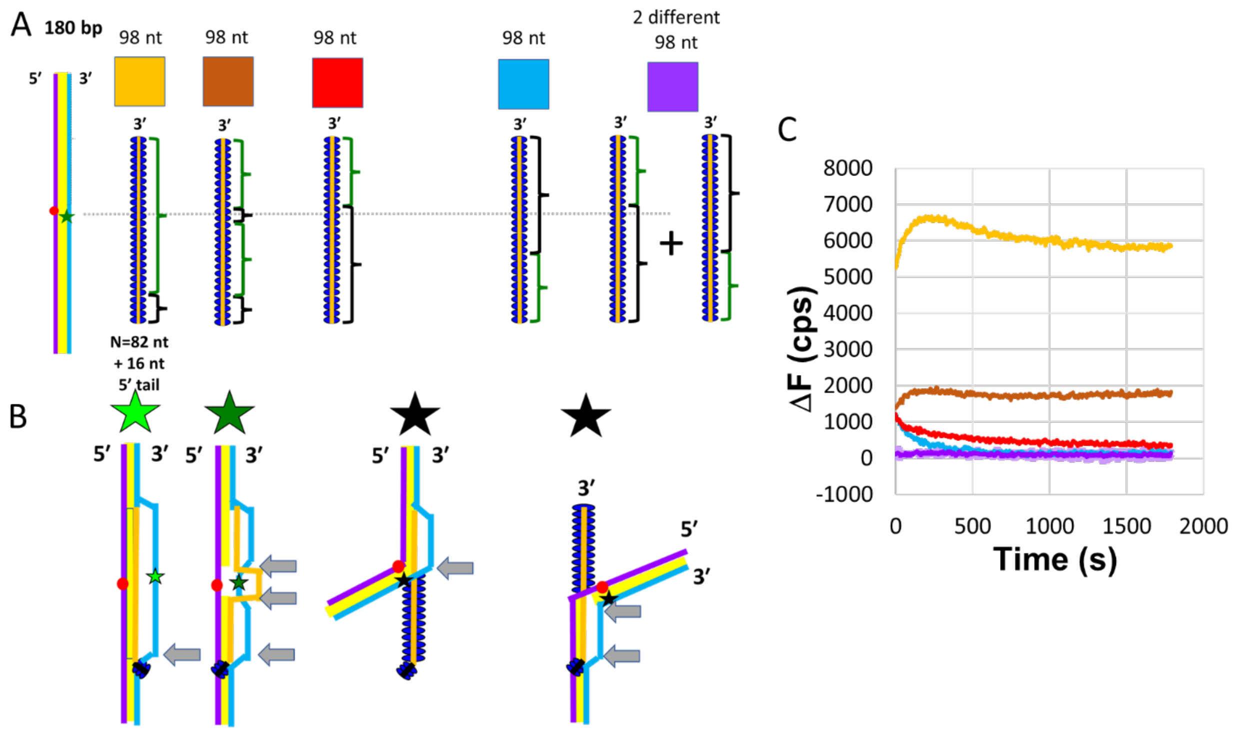


Figure 3 — figure supplement **1 Strand exchange across two mismatched triplets monitored using FRET.** A. Schematic of interactions. The purple, light blue, and orange lines represent the complementary, outgoing, and invading strands. The dark blue ovals represent RecA. The red circle and green star show the locations of the fluorophores along the 180 bp dsDNA. All of the invading strands contain 98 nt. The 16 nt nearest the 5′ end are always heterologous to the dsDNA, as indicated by the black brackets that highlight the heterologous regions. The homologous regions are highlighted by the magenta brackets. The boundaries between homologous and heterologous regions are indicated by the gray arrows. The first filament on the left includes 82 contiguous bp that are homologous to the dsDNA. The second filament includes 47 homologous bases at the 3′ end, interrupted by a 6 bp region of heterology, followed by a 29 bp homologous region. The third filament includes 47 homologous bp starting at the 3′ end of the filament. The fourth filament includes a 29 bp region of homology adjacent to the 16 nt heterologous tail at the 5′ end. The fifth case is the simultaneous addition of the third and fourth filaments to the dsDNA. B. Illustrates strand exchange products that could be formed by combining the dsDNA with the filament shown directly above the products. The color of the star indicates the expected fluorescein emission for the strand exchange product shown. Bright green, medium green, and black indicate high emission, some emission, and strong quenching, respectively. C. Change in fluorescein emission (ΔF) vs time curves for the various cases, the color of the curve corresponds to the color of the box above the schematic. Thus, the orange and dark orange curves correspond to a filament with 82 homologous nt and a filament with a 6 nt heterologous region surrounded by two homologous regions. The red and light blue curves show results for filaments that are homologous to either the 3′ or 5′ side of the 6 heterologous bp in the second filament. The purple curve represents the results when both of those filaments are added to the dsDNA simultaneously. Error bars on the purple curve are shown in lavender. Only the filament with 82 uninterrupted homologous bases and the filament with homology on both sides of the 6 heterologous bases creates a significant FRET signal, suggesting that strand exchange can progress through six contiguous heterologous bp and form a long-lived strand exchange product if and only if the heterology is flanked by a significant region of homology.
