## Supplemental Data 7 for "RecA balances genomic stability and evolution using many successive mismatch tolerant homology tests"

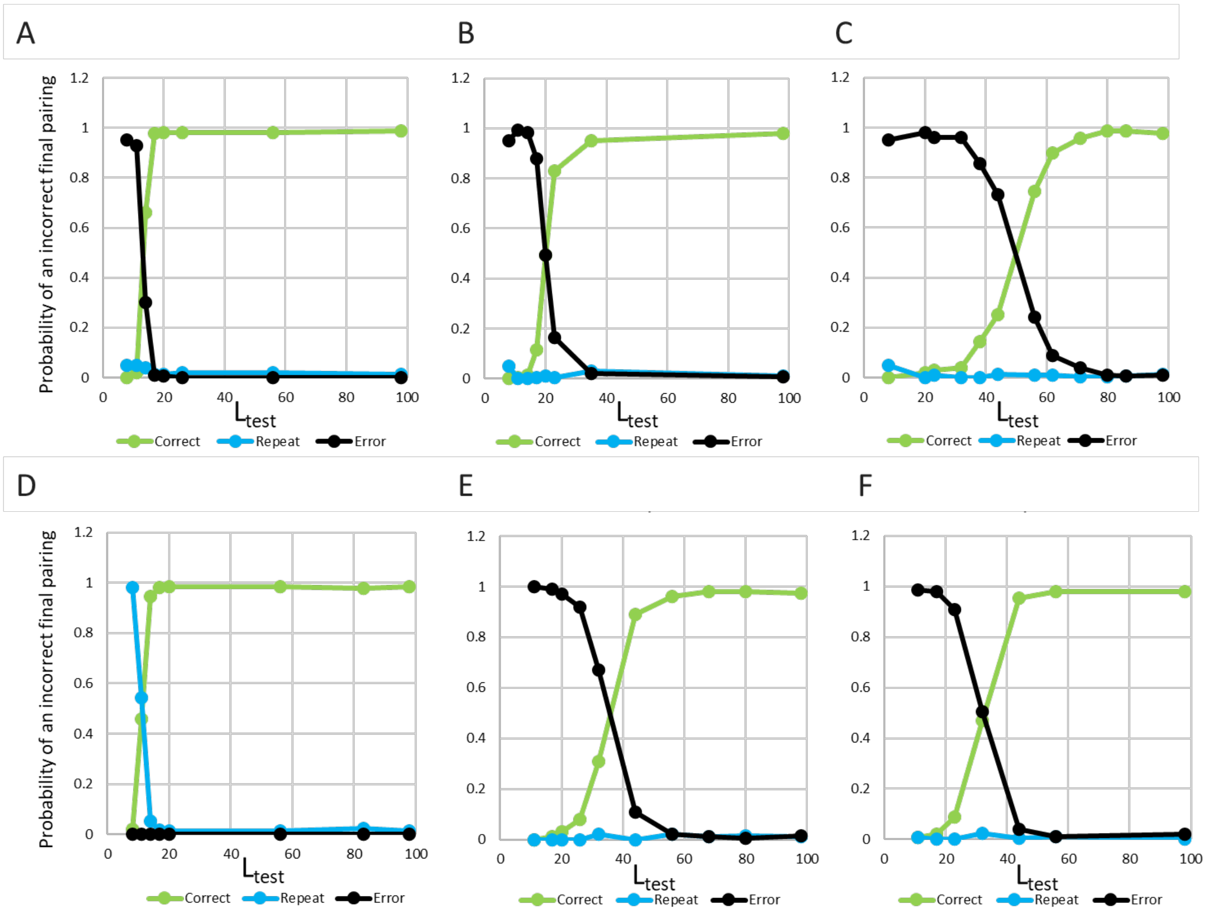


Figure 2 — figure supplement 6 **Characterization of irreversible pairings for the simplified model applied to the *E. coli* MG1655 genome.** Results are divided into correct pairings (green), pairings that contain no mismatches but join different copies of exact repeats (blue), and pairings that contain mismatches (black) as a function of $L_{test}$. A. Results for $N_{mismatch}$ = 0. B. Results for $N_{mismatch}$ = 1. C. Results for $N_{mismatch}$ = 2. D. Results for homology testing that accepts no mismatches at all. E. Results for the probabilistic model of triplet testing with 100%, 75%, 50%, and 25% probability of passing with 0, 1, 2, and 3 mismatches, respectively. F. Results for the probabilistic model with the same triplet probabilities as D, but the 8 bp test does not accept more than one mismatch and has only 25% probability of accepting one mismatch.
