## Supplemental Data 8 for "RecA balances genomic stability and evolution using many successive mismatch tolerant homology tests"

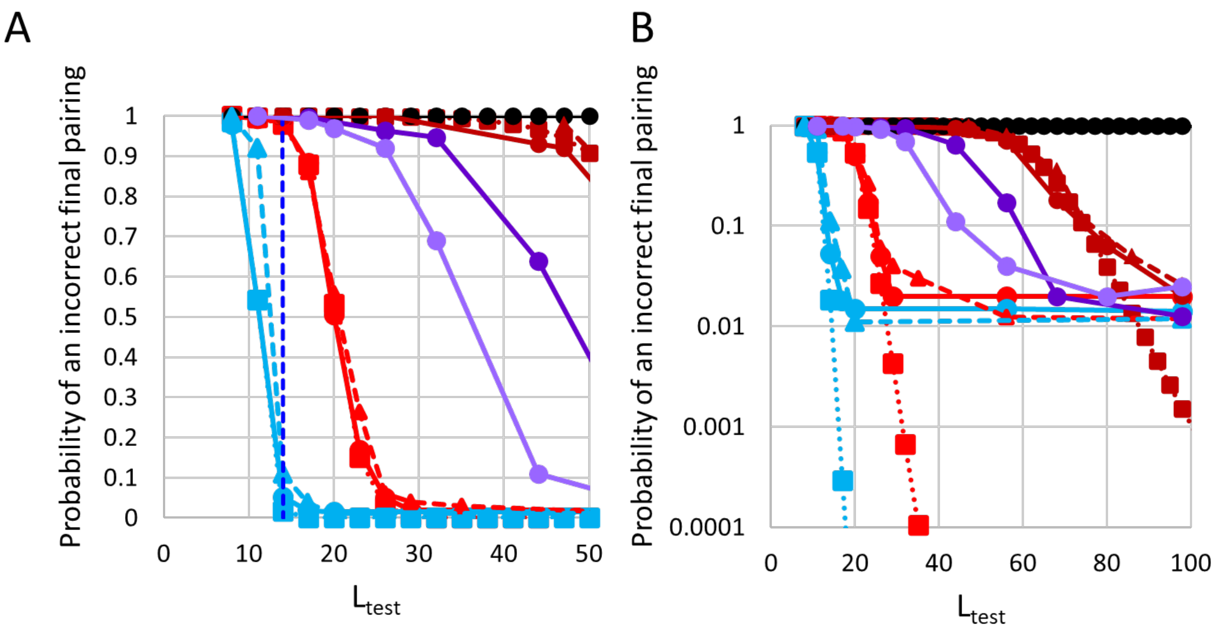


Figure 2 — figure supplement 5 **Probability that a final product is incorrect vs. $L_{test}$ when the 8 bp test and the triplet tests accept the same number of mismatches when applied to the *E.coli* MG1655 genome.** The blue, bright red, dark red, and black curves show results for $N_{mismatch}$ = 0, 1, 2, and 3 respectively. Dashed Curves with triangle symbols represent results for invading strands that have Chi sites on the 3′ ends. Results are similar for any termination near the Chi sites. The light purple curves indicate results for the probabilistic model of triplet testing with 100%, 75%, 50%, and 25% probability of passing 0, 1, 2, and 3 mismatches, respectively. The dark purple curves indicate results for the probabilistic model of triplet testing with 100%, 5%, 50%, and 50% probability of passing 0, 1, 2, and 3 mismatches, respectively.
